# Bifidobacteria-mediated immune system imprinting early in life

**DOI:** 10.1101/2020.10.24.353250

**Authors:** Bethany M. Henrick, Lucie Rodriguez, Tadepally Lakshmikanth, Christian Pou, Ewa Henckel, Axel Olin, Jun Wang, Jaromir Mikes, Ziyang Tan, Yang Chen, Amy M. Ehrlich, Anna Karin Bernhardsson, Constantin Habimana Mugabo, Ylva Ambrosiani, Anna Gustafsson, Stephanie Chew, Heather K. Brown, Johann Prambs, Kajsa Bohlin, Ryan D. Mitchell, Mark A. Underwood, Jennifer T. Smilowitz, J. Bruce German, Steven A. Frese, Petter Brodin

## Abstract

Immune-microbe interactions early in life influence an individual’s risk of developing allergies, asthma and some autoimmune disorders. Breastfeeding helps guide the development of healthy immune-microbe relationships, in part by providing nutrients to specialized microbes that in turn benefit the host and its developing immune system. Such bacteria having co-evolved with humans are associated with reduced risks of immune mediated diseases but are increasingly rare in modern societies. Here we map an immunological sequence of events, triggered by microbial colonization that distinguish children with different gut bacterial composition. Lack of bifidobacterial species is associated with elevated markers of intestinal inflammation and immune dysregulation and in a randomized trial of breastfed infants, the infant-adapted *Bifidobacterium infantis* EVC001 silenced intestinal Th2 and Th17 immune responses, while inducing IFNβ, and its metabolites skew T-cell polarization *in vitro*, from Th2 towards Th1, suggesting a healthier immune imprinting during the first critical months of life.

**HIGHLIGHTS:** An ordered sequence of immune changes after birth, driven by microbial interactions

Low gut *Bifidobacterium* abundance is associated with markers of intestinal inflammation

Feeding *B. infantis* EVC001 silenced intestinal Th2 and Th17 but upregulates IFNβ

*B. infantis* EVC001 metabolites and/or enteric cytokines skew naïve T-cell polarization towards Th1 and away from Th2

## INTRODUCTION

Mounting evidence indicate that the composition of the infant gut microbiome is critical to immunological development, particularly during the first 100 days of life in which aberrations in gut microbial composition are most influential in impacting the developing immune system. Indeed, multiple studies have emphasized how early gut microbiome dysbiosis, described as an overabundance of Proteobacteria (Shin et al., 2015) and loss of ecosystem function (Duar et al., 2020) is associated with both acute and chronic immune dysregulation, leading to common conditions such as colic (Rhoads et al., 2018), atopic wheeze and allergy (Arrieta et al., 2015; Laforest-Lapointe and Arrieta, 2017) and less common, but serious immune mediated disorders such as type-1 diabetes (Vatanen et al., 2016) and Crohn’s disease (Hviid et al., 2011). Recent developments in systems immunology now enable us to profile immune system development in humans at the systems level, and map cell-cell regulatory relationships in health and disease (Davis and Brodin, 2018). Further, small blood volumes available from newborn children are no longer prohibitive and as little as 100μL of whole blood is sufficient for systems-level immunomonitoring (Brodin et al., 2019). Previously, in an analysis of immune development in human newborns, we identified drastic modulation in immune cell and protein abundances that were largely shared among preterm and term children, pointing towards a stereotypic postnatal adaptive response to environmental exposures (Olin et al., 2018). Moreover, gut dysbiosis in a subset of these infants was associated with elevated markers of immune activation in the blood (Olin et al., 2018).

Dysbiosis of the infant gut microbiome is common in modern societies and a likely contributing factor to the increased incidences of immune mediated disorders (Dominguez-Bello et al., 2019; Mohammadkhah et al., 2018; Sonnenburg and Sonnenburg, 2019). Therefore, there is great interest in identifying microbial factors that can support healthier immune system development and potentially prevent some cases of allergy, autoimmunity and possibly other conditions involving the immune system (Renz and Skevaki, 2020). Loss of *Bifidobacterium* early in life is one commonly reported factor in infants more likely to develop autoimmune activity as seen in a birth cohort in Finland (Vatanen et al., 2016) and atopic wheeze in another cohort in rural Ecuador (Arrieta et al., 2017). Moreover, observational studies have identified a link between the loss of *Bifidobacterium* in infants and enteric inflammation early in life (Henrick et al., 2019; Rhoads et al., 2018).

Human breastmilk contains abundant Human Milk Oligosaccharides (HMOs) and these are not digestible by humans who lack necessary glucosidases (Sela and Mills, 2010). Instead, the maternal energy used to create these complex sugars is justified by the importance of providing a selective nutritional advantage to microbes specialized in metabolizing HMOs and serving an evolutionary important role in the gut of newborn infants. *Bifidobacterium longum subspecies infantis* (*B. infantis*) is one example of a bacterial strain adapted to metabolizing HMOs (LoCascio et al., 2010; Sela et al., 2008; Underwood et al., 2015). *B. infantis* is commonly found in breastfed infants in countries with low incidence of immune mediated disorders, such as Bangladesh (Huda et al., 2014), and Malawi (Grzekowiak et al., 2012), but rare in Europe (Abrahamsson et al., 2014; Avershina et al., 2014; Jost et al., 2012; Roos et al., 2013) and in North America (Azad et al., 2013; Lewis et al., 2015). Strategies for introducing *B. infantis* have been successfully accomplished, such as EVC001 (Evolve BioSystems Inc.), which is able to stably and persistently colonize and achieve a dominant composition in the intestinal microbiome of breastfed infants leading to reduced markers of intestinal inflammation such as fecal calprotectin (Henrick et al., 2019).

Here, we extend previous findings by combining systems-level analyses of immune development and microbiome colonization in 155 Swedish infants and find that low abundance of gut bifidobacteria is associated with intestinal inflammation and immune dysregulation at three months of life. We also demonstrate a silencing of intestinal inflammation in infants supplemented with *B. infantis* EVC001 (Evolve BioSystems Inc.) in an intervention study in California. Fecal water from infants colonized by *B. infantis* EVC001 skew naïve T-cell polarization *in vitro* from Th2 towards Th1, and modulate Th17 cell states, indicative of a healthier immune system imprinting during the first critical months of life.

## RESULTS

### Sequential waves of immune cell expansions during the first months of life

We analyzed longitudinal blood samples (n=406) postnatally from infants (n=155) born at the Karolinska University Hospital between April 2014 to December 2019 using a mass cytometry panel with 44 antibodies targeting activation and differentiation markers across all immune cell populations. In total, we profiled 110 immune cell populations from 270 blood samples and quantified 267 unique plasma proteins using using Olink assays (Lundberg et al., 2011) (Olink, Uppsala, Sweden), which elucidated immune responses and developmental changes (Olin et al., 2018) as well as infectious disease susceptibility postnatally (Pou et al., 2019). We also collected paired fecal samples (n=168) performed shotgun metagenomic sequencing from these same children and timepoints (Figure 1A).

**Figure 1.**
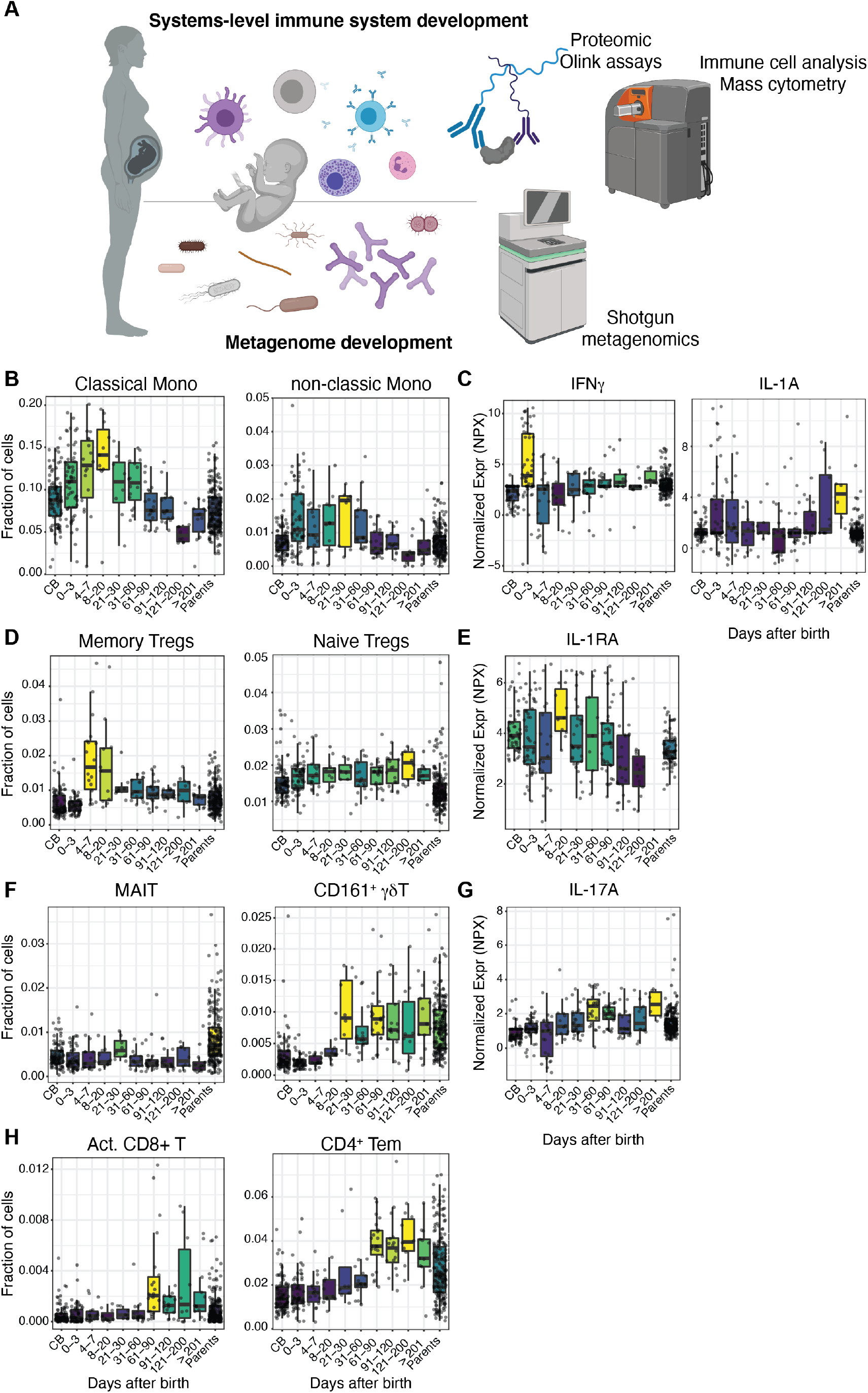
Systems-level analysis of immune development in human newborns. **(A)** Study outline, **(B)** Monocyte abundance analyzed by Mass cytometry from blood samples (n=4O6) binned by sampling day of life. Box plots colored by mean rank. (C) Blood IFNγ and IL-1A analyzed by Olink assays from plasma samples (n=4O6) binned by sampling day of life and box plots colored by mean rank. Mass cytometry analyses of **(D)** Tregs, **(E)** IL-1RA, **(F)** MAIT and CD161-expressing γδT-cells, **(G)** IL-17A, **(H)** activated CD8^+^ T-cells (HLA-DR÷ CD38^+^) and Effector memory CD4^+^ T-cells defined by CD27 and CD45RA expression.

When ordering immune cell frequency measurements by day of sampling, we observed an early expansion of classical and non-classical monocytes during the first few days of life (Figure 1B), and a concomitant surge in circulating IFNγ, and a less pronounced increase in IL-1α (Figure 1C). Starting a few days after this monocyte expansion, a significant increase in memory Tregs was observed uniformly across children, while naïve Tregs abundance remained stable (Figure 1D). This expansion of antigen experienced Tregs indicate a tolerizing response to antigen following the initial innate reaction mediated by monocytes. This memory Treg expansion is also associated with a transient increase in circulating IL-1RA at day 21-30 (Figure 1E), further emphasizing a balanced response to initial innate reactions starting immediately after birth. Gut associated cells such as MAIT cell populations were found at a consistently lower frequency as compared to parents, but with a small transient increase around 1 month of life (Figure 1F).

At this same timepoint a more robust increase in circulating CD161^+^ γδT cells was observed and reached cell frequencies similar to adult levels (Figure 1F). Such cells are important producers of IL-17A (Maggi et al., 2010), which was corroborated by increased levels of IL-17A in plasma during the same initial 3-month time frame (Figure 1G). A final activation event involve the expansion of CD4^+^ and activated CD8^+^ αβT-cells from 2 months onwards (Figure 1H), and collectively these findings indicate transient immune activation events in early and mid-infancy, likely triggered by colonizing microbes like the weaning reaction described in mice (Nabhani et al., 2019), but noticeably different with respect to timing and immune cell populations involved.

### Expansion of mucosal-specific CD4^+^ T-cells in the blood of newborn children

Work in murine model systems and human subjects have revealed that immune cells primed by antigens at mucosal surfaces circulate and are detectable in the peripheral blood. Specifically, memory CD4^+^ T-cells identified by the expression of CD38 and lacking the lymphoid tissue homing marker CD62L are examples of such mucosal-specific T-cells that are abundant in the intestinal tissue but also identifiable in the blood of humans at a frequency of ~4-8% of total CD4^+^ T-cells (Pré et al., 2011). Here, we identified the same population of CD4^+^ T-cells in the blood of newborn children; however, at a greater abundance of 33.6% and expanding in the blood during the first weeks of life to reach more than 60% of memory CD4^+^ T-cells within the first 100 days of life (Figure 2A). Since these cells have downregulated CD45RA we believe these cells have responded to their native antigen, presumably at the mucosal surfaces and undergone memory phenotype transition. To understand the functional aspects of these T-cells, we sorted these by flow cytometry and performed bulk mRNA-sequencing. We also sorted out total memory CD4^+^ T-cells from the same samples and compared blood-transcriptional modules (BTMs)(Li et al., 2013) between mucosal-specific memory CD4^+^ T-cells and total memory CD4^+^ T-cells. The most enriched hallmark pathways in mucosal-specific memory CD4^+^ T-cells were type-I and type-II IFN responses (Figure 2B) and the most upregulated genes in mucosal specific memory CD4^+^ T-cells included the Complement regulatory Factor H (CFH), the cytokine IL-15 important for NK cell homeostasis as well as the monocyte growth factor CSF1 (M-CSF), suggesting that the mucosal-specific memory CD4^+^ T-cells serving a regulatory function vis-à-vis NK-cells and monocytes in the newborn intestinal mucosa (Figure 2C-D).

**Figure 2.**
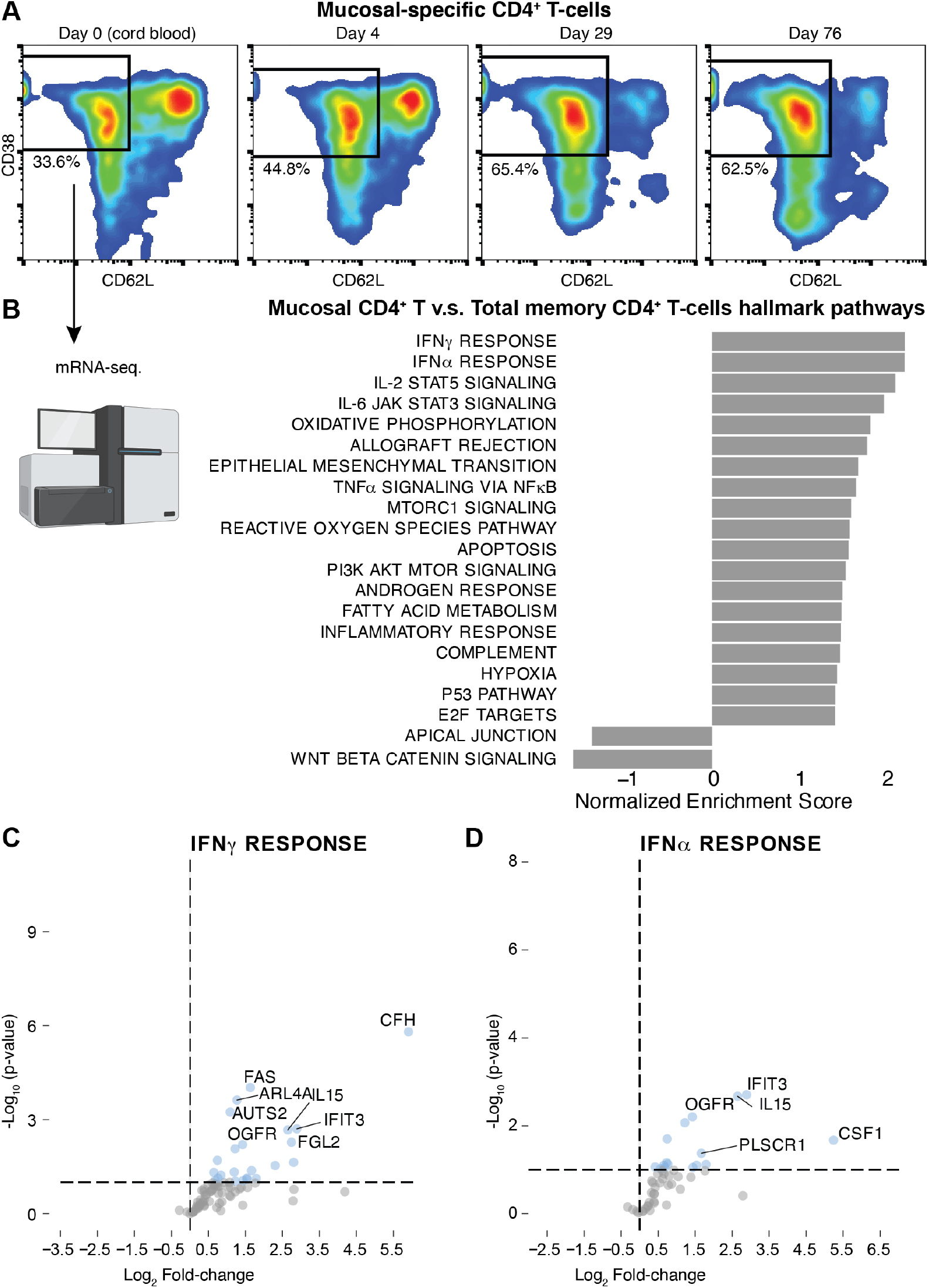
Mucosal specific CD4^+^ T-cells expand and perform type-1 and II IFN-responses. **(A)** Representative FACS plots of CD38^+^ CD62L-CD4^+^ T-cells collected at postnatal day 0, 4, 29 and 76. **(B)** Bulk mRNA-seq. and gene set enrichment analysis showing top enriched hallmark pathways in mucosal-specific v.s. total memory CD4^+^ T-cells. Volcano plots of differentially regulated mRNA transcripts in mucosa-specific CD4^+^ T-cells vs total memory CD4^+^ T-cells within the hallmark pathways **(C)** IFNγ responses and, **(D)** IFNα responses.

### Variable colonization of the infant gut after birth

To better understand the expansion of mucosal-specific memory CD4^+^ T-cells and the microbial antigens potentially driving the observed response early after birth, we performed shotgun metagenomic sequencing of longitudinal fecal samples from the same children. We focused on bacterial components and found a highly variable alpha-diversity at birth but also a trend towards increasing alpha-diversity over time after birth (Figure 3A). The most variable genera over time were identified partial least square regression analysis vs. days after birth and a latent vector explaining changes over time extracted. The top contributing genera are shown as fraction of reads vs. time after birth and colored by mode of delivery (Figure 3B). Several notable changes are seen, and largely in line with previous literature such as high abundance of *Bacteroides* in vaginally delivered children while staphylococci and streptococci are abundant in cesarean-delivered infants (Shao et al., 2019) (Figure 3B).

**Figure 3.**
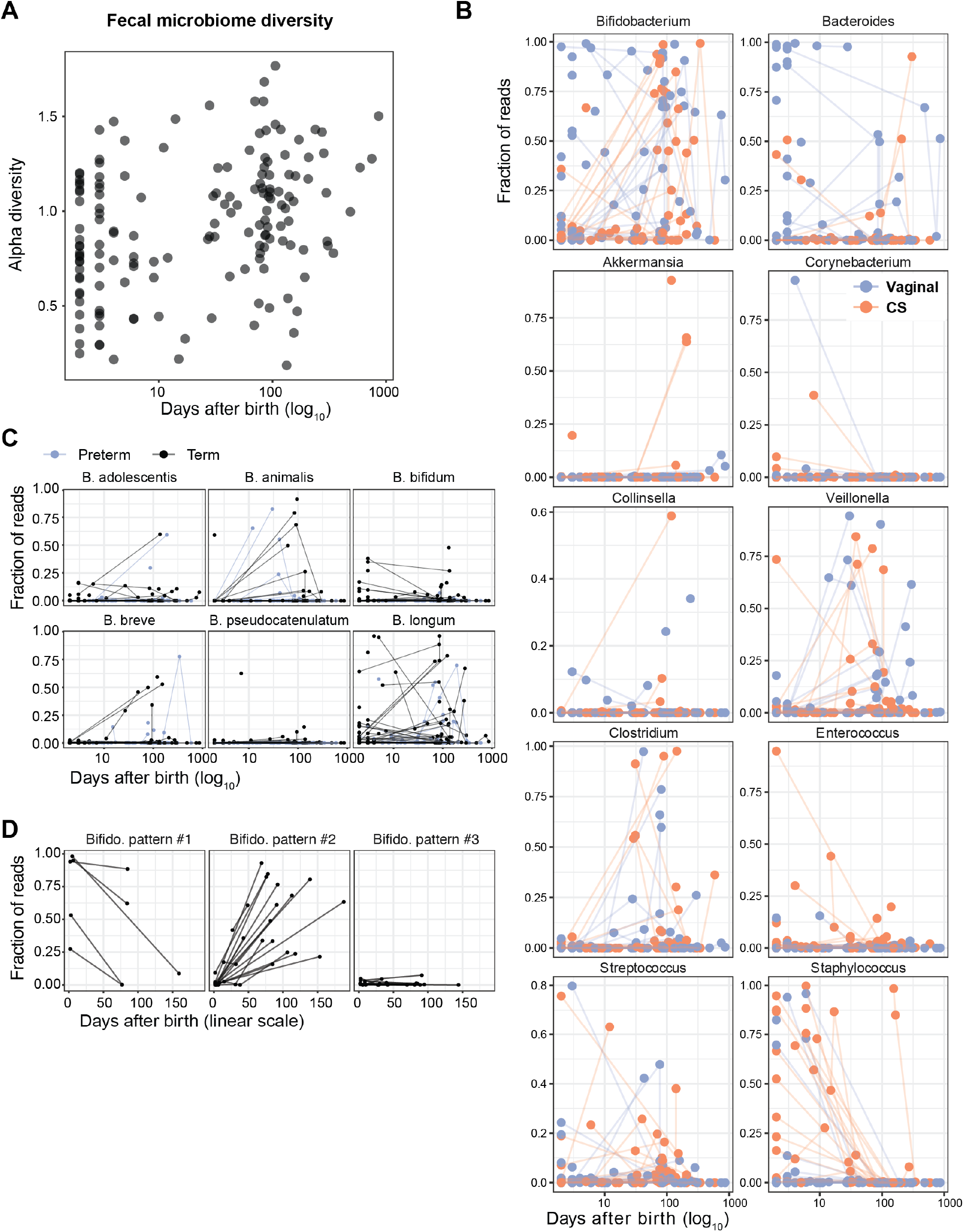
Fecal microbiome development. **(A)** Alpha diversity of n=?68 fecal samples collected longitudinaly from infants. **(B)** Bacterial genera changing most over time after birth. **(C)** Species level Bifidobacterial abundances. Only species detected in at leat one sample shown. **(D)** Three principal pattern of bifidobacterial colonization among children in our cohort.

There is a drastic expansion of Bifidobacteria occurring after birth in the majority of newborns, and in some infants, all identified reads are mapping to the *Bifidobacterium* genus (Figure 3B), indicating their potential to dominate the gut microbiome early in life, particularly in breastfed infants. We found that multiple species of bifidobacteria expanded early in life in both preterm and term children, in particular *B. longum, B. breve*, and *B. animalis* (Figure 3C). Given these findings, the changes in bifidobacteria can be used as a surrogate markers of gut microbiome evolution in the newborn children with three principal patterns seen in our cohort, 1) rare cases of abundant bifidobacteria that are lost over time, 2) the most common pattern of increased abundance of bifidobacteria over time, and 3) children with very rare, or even undetectable levels of bifidobacteria over time during the first three months of life (Figure 3D). Given the known associations between Bifidobacterial abundance and the risk of developing immune mediated diseases in the future (Arrieta et al., 2015, 2017; Vatanen et al., 2016) we were interested to explore immune system development in children with these different surrogate patterns of microbial colonization.

### Immune dysregulation in infants not expanding gut Bifidobacteria

To capture imprinting effects, we focused on blood immune measurements around three months of life involving 268 plasma proteins and all major immune cell populations in blood. We compared these immune system features between infants with (pattern #2) and without bifidobacterial expansion (Pattern #3). We find that 3-month immune cell composition in infants with low gut bifidobacterial abundance was characterized by expanded populations of effector cells and proinflammatory immune cell populations such as proinflammatory monocytes and MAIT-cells, important T-cells in the intestinal immune system compartment and known to respond to bacterial Vitamin B-metabolites in the mucosa (Ioannidis et al., 2020) (Figure 4A).

**Figure 4.**
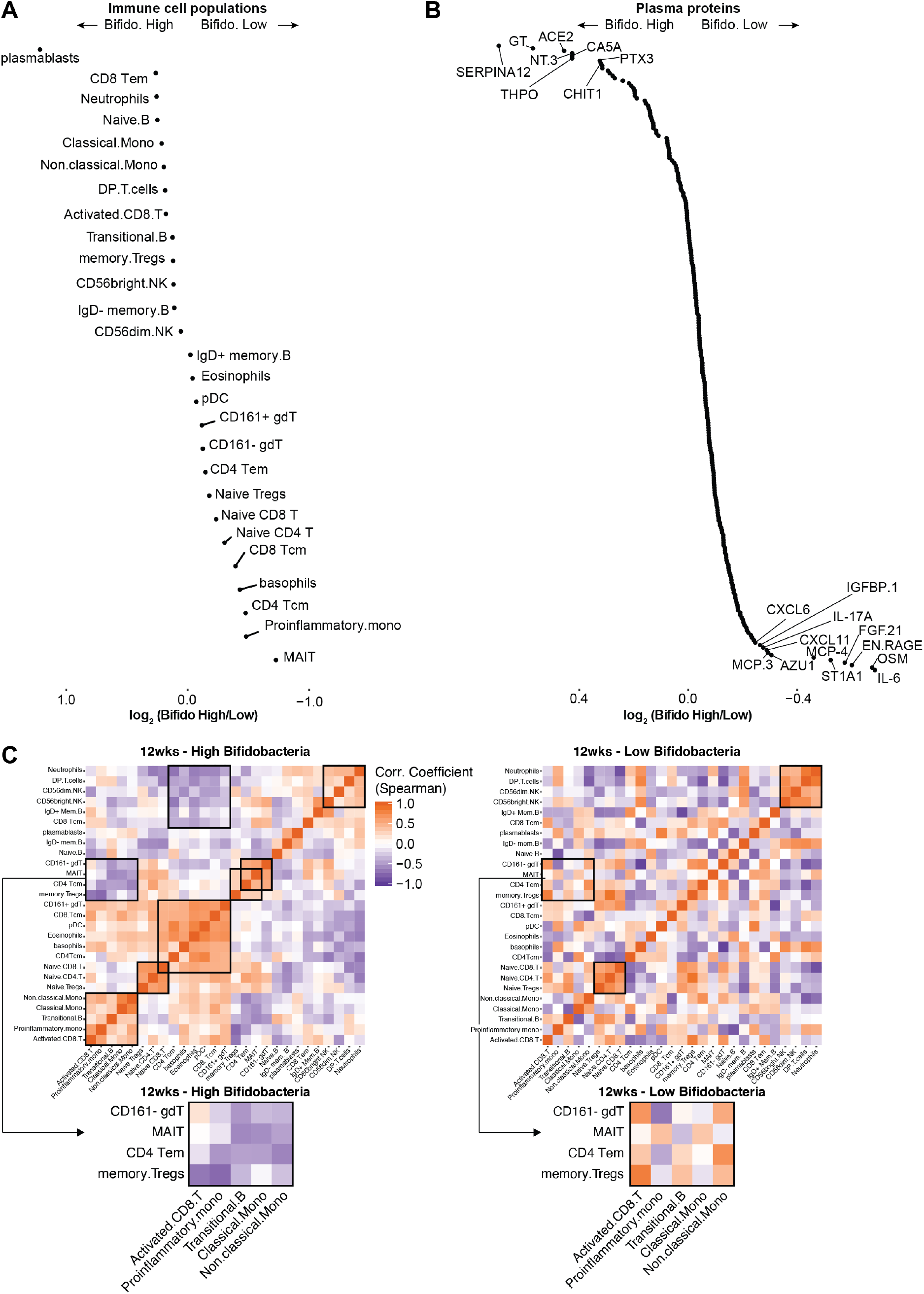
Immune dysregulation in infants with low v.s high bifidobacteria. **(A)** Fold-change immune cell frequencies at 3 months of life in infants with high v.s. low fecal bifidobacteria. **(B)** Fold-change plasma protein levels at 3 months of life in infants with high v.s. low fecal bifidobacteria. **(C)** Spearman correlation matricies of immune cell frequencies at 3 months of life in children with high v.s. low fecal bifidobacteria. Black boxes highlight modules of co-regulated immune cell populations.

In contrast, children with abundant gut Bifidobacteria have elevated plasmablast frequencies and also higher frequencies of antigen-experienced regulatory T-cells which are both essential for healthy immune-microbe relationships in the intestine (Smith et al., 2013). The plasma protein analyses revealed similarly upregulated markers of inflammation such as IL-6 and the Oncostatin M (Figure 4A). The latter is notable given results from West et al, implicating the latter protein as a mediator of intestinal inflammation in patients with inflammatory bowel disease (West et al., 2017). To better understand the regulatory relationships in the newborn immune system in these groups of infants, we assessed the cell-cell dependencies. To this end, we calculated Spearman correlation matrices in infants with high levels of *Bifidobacterium* versus low *Bifidobacterium* abundance, respectively (Figure 4C). Such cell-cell correlation matrices can be informative by revealing coregulated cell populations, and in particular, context-dependent differences to this co-regulation network (Rodriguez et al., 2020). We found that selected cell-cell relationships are shared among high levels of *Bifidobacterium* versus low *Bifidobacterium* abundance including strong positive correlation among naïve CD4^+^, naïve CD8^+^ and naïve regulatory T-cells, likely reflecting the overall thymic output of an individual child (Figure 4C). Other modules of coregulated cell populations were markedly different between the groups of children.

In particular, in infants with higher *Bifidobacterium* abundance, we found that memory-phenotype regulatory T-cells are inversely correlated with the levels of activated CD8^+^ T-cells, proinflammatory monocytes and few additional immune cell populations, whereas this relationship is completely abolished in infants at three months of life, who have low abundance of *Bifidobacterium* (Figure 4C). In contrast, these children have strong positively correlated levels of memory Tregs and activated CD8^+^ T-cells, suggesting ongoing immune activation, rather a homeostatic balance between tolerance and inflammation (Figure 4C). We conclude that infants who are not colonized by bifidobacterial species during their first months of life have an immune system state at three months of life characterized by elevated markers of intestinal inflammation, activated immune cell populations and a perturbed immune cell network.

### *B. infantis* EVC001 supplementation silenced intestinal inflammation in newborn children

In an attempt to improve immune system education and silence the chronic inflammatory responses seen in children devoid of infant-adapted bifidobacterial species, we evaluated fecal cytokines in exclusively breastfed, term infants, half of whom were fed *B. infantis* EVC001 daily from day 7 to day 28 postnatal and half that were not fed a probiotic. Baseline fecal cytokine levels were not significantly different between the two groups; however, infants fed *B. infantis* EVC001 had significantly lower levels of Th2-inducing cytokine, IL-4 and IL-13, proinflammatory cytokine, IL-17A, and regulatory and chemotactic cytokines, IL-21, IL-31, IL-33, and MIP3a by day 60 postnatal (*P*=0.022, 0.009, 0.046, 0.0002, 0.0038, 0.00001, and 0.0038, respectively; Figure 5B). Conversely, IFNβ concentrations were significantly increased in infants fed *B. infantis* EVC001 (*P*=0.017; Figure 5B). This suggested a modulatory effect of proinflammatory and Th2 responses by *B. infantis* EVC001 colonization, so we performed pairwise correlation tests for all 40 individual stool samples at day 60 of life comparing microbial composition and fecal cytokine concentrations (Spearman correlation with Benjamini-Hochberg false discovery rate correction a < 0.02), which identified three taxa, including *Clostridiaceae, Enterobacteriaceae*, and *Staphylococceae* significantly correlated with proinflammatory cytokine production. Specifically, *Clostridiaceae* abundance correlated with increased IL-21, IL-33, and IL-4, *Enterobacteriaceae* correlated with increased IL-13, IL-17A, IL-21, IL-33, and IL-4 production, while *Staphylococceae* correlated with higher levels of IL-21 (Figure 5C). Additionally, we used a Random Forest analysis to identify taxa associated with increased IFNβ production and predictors with the 10 lowest mean depth values confirming that *Bifidobacteriaceae, Streptococcaceae, Veillonellaceae*, and *Staphylococcaceae* most often contributed to IFNβ concentration (Supplementary Figure 1A).

**Figure 5.**
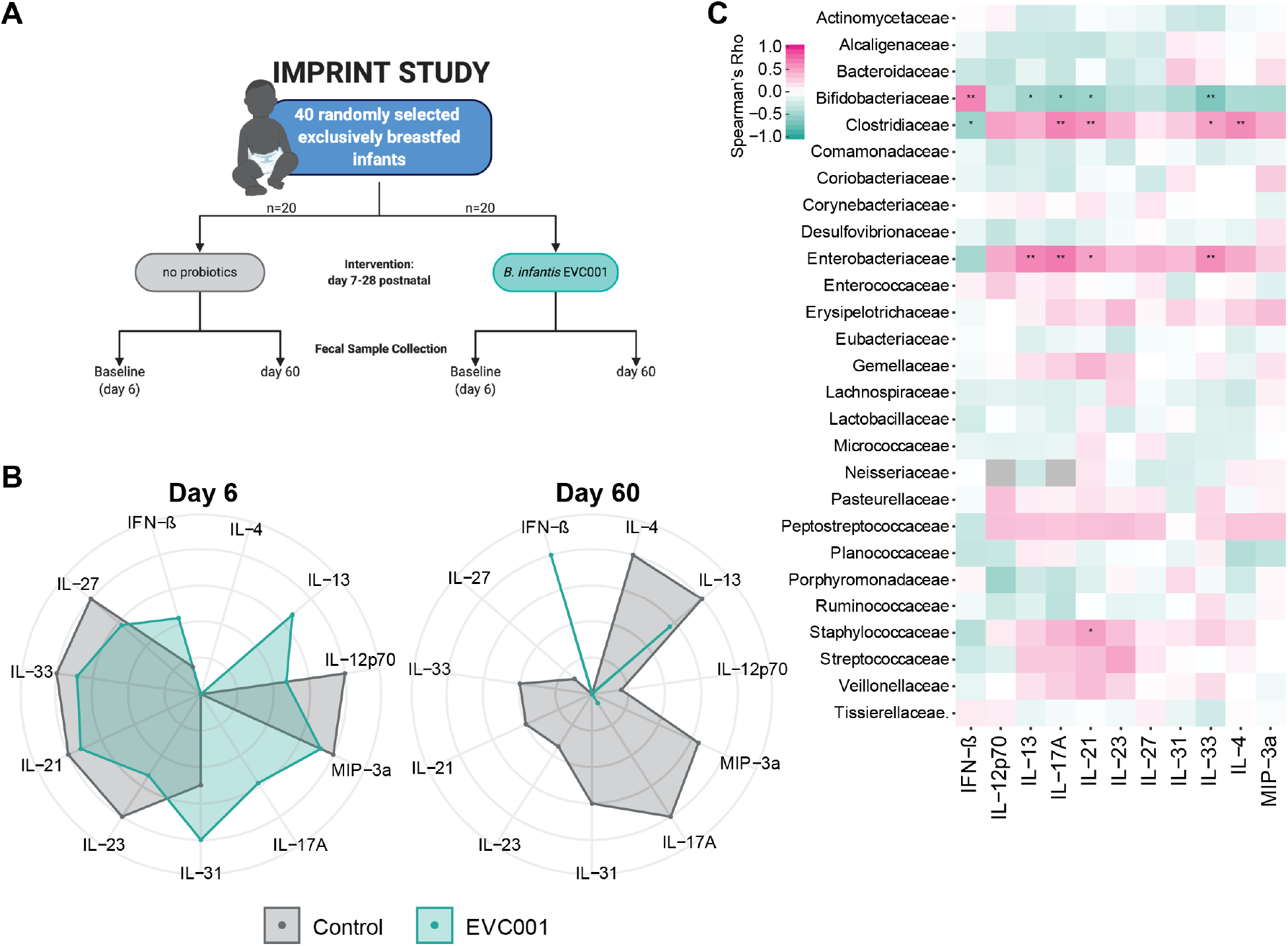
B. infantis EVC001 supplementation is associated with reduced intestinal inflammation. **(A)** IMPRINT study design and randomization, **(B)** Fecal cytokines at baseline (Day 6) and post-treatment with B.infantis EVC001, or no supplementation. **(C)** Spearman correlation coefficients between fecal cytokine levels and bacterial abundance. *P=0.05, **p=0.01.

We next evaluated whether IFNβ concentration at day 60 correlated with early colonization of *Bifidobacteriaceae* at day 21 postnatal. Fecal samples containing *Bifidobacterium* at day 21 had significantly increased levels of IFNβ compared to samples that did not contain *Bifidobacterium* by day 60 postnatal (Supplementary Figure 1B), and at a the species level, *B.longum* abundance at day 21 correlated with IFNβ levels at day 60 postnatal (Supplementary Figure 1C; *P* = 0.0028, rho = 0.48). Importantly, no other *Bifidobacterium* species correlated with increased IFNβ levels indicating that *B. longum* abundance may drive increased IFNβ production. These findings indicate that supplementation with a Bifidobacterial strain, adapted to expand in breastfed infants, may silence proinflammatory responses in the gut and potentially induce a healthier immune system priming.

### *B. infantis* EVC001 metabolites influence T-cell polarization

Classically, it is suggested that newborn infants have an immune system biased towards Th2 responses rather than Th1 proinflammatory responses, yet this is mostly based on data from cord blood cells collected prior to any postnatal environmental exposures, or mouse model systems (Brodin, 2020). Our evaluation of enteric cytokines showed significantly decreased Th2 cytokines, IL-4 and IL-13, Th17 cytokine IL-17A, as well as increased IFNβ concentrations in infants colonized with *B. infantis* EVC001 compared to those that were not; therefore, we investigated the potential for microbial metabolites and enteric cytokines from two infant populations that were exclusively fed human milk but were discordant for *B. infantis* colonization to skew T-cell polarization in vitro. We flow-sorted naïve CD4^+^ T-cells from a healthy adult donor and polarized these cells using standard cytokine combinations (Cano-Gamez et al., 2020) in combination with fecal waters (1:100 dilution) collected from newborns fed *B. infantis* EVC001 or control children lacking *B. infantis* (Figure 6A). To evaluate the polarization and cellular states of the T-cells in the cultures we applied a targeted multiomics approach (Rhapsody, BD Biosciences) in which we combined the analysis of 259 mRNA by targeted sc-mRNA-sequencing and 10 proteins using oligo-coupled antibodies (BD Abseq™, BD Biosciences)(Mair et al., 2020). In this way we were able to assess transcription factor and cytokine gene expression often more difficult to detect in untargeted sc-mRNA-sequencing applications, and although the overall UMAP distribution of cells is similar across conditions, subtle differences in cell states were observed (Figure 6B). Using a graph abstraction and clustering method, PAGA (Wolf et al., 2019), we found that T-cell states differ by polarizing condition as expected, but we also observed differences between cells cultured in the presence of *B. infantis* EVC001 fecal water compared to fecal water lacking *B. infantis* (Figure 6C). Specifically, Th1, Th2 and iTregs are largely comparable between *B. infantis* EVC001 and no *B. infantis* fecal water (Figure 6C). Conversely, Th0 cells cultured without additional polarizing cytokines and no *B. infantis* fecal water induced a cell state similar to that of Th2-polarized cells, while the *B. infantis* EVC001 induced a Th0 cell state that more closely reflect Th1-induced conditions (Figure 6C). Furthermore, we assessed the differentially regulated genes among cultured Th0 cells exposed to fecal waters from the two different groups and found upregulation of GZMA, GZMB, TNF and STAT1 in the cells skewed towards a Th1-phenotype after exposure to *B. infantis* EVC001 fecal water while IL23R expression in the cells exposed to no *B. infantis* fecal water, similar to the Th2-like cells (Figure 6D). These findings suggest *B. infantis* EVC001 specific metabolites and/or enteric cytokines induced in the intestine by the presence of *B. infantis* EVC001 polarizes naïve CD4^+^ T-cells towards the Th1 lineage, and are of particular interest given the association between Th2-type cells and the development of immune-mediated diseases such as asthma and allergies in young children in which a polarizing effect away from Th2 towards Th1 might prove beneficial in the future.

**Figure 6.**
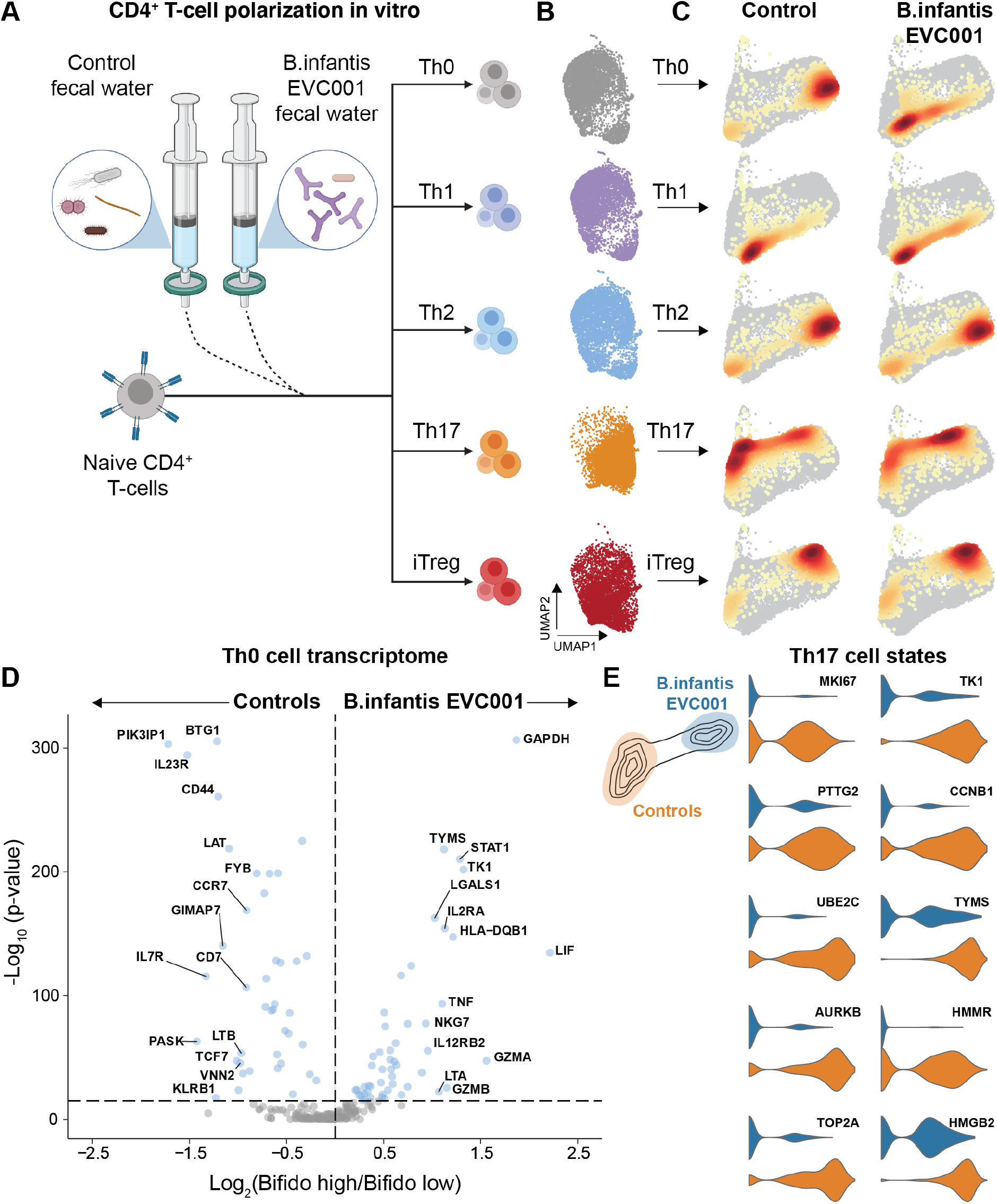
CD4^+^ T-cell polarization under the influence of microbial metabolites. **(A)** CD4^+^ T-cell polarization in vitro in the presence of fecal water from infants supplemented with B.infantis EVC001 or control (No supplement). **(B)** UMAP plots of polarized T-cells analyzed by targeted sc-mRNA-seq. **(C)** PAGA plots of T-cells polarized in the presence of fecal water from fecal water from infants given B.infantis EVC001 supplementation or control. Coloring by cell density from grey (low) to red (high). **(D)** Volcano plot showing differentially expressed mRNA in ThO cells cultured with fecal water from infants given B.infantis EVC001 supplementation or control. **(E)** Top 10 differentially expressed mRNA in Th17 cell states induced by fecal water from infants given B.infantis EVC001 supplementation or control.

Apart from the skewing of Th0 cells by fecal water, we also saw a slight difference in the cell states among Th17-polarized cells when comparing *B. infantis* EVC001 and no *B. infantis* fecal water effects (Figure 6C). Specifically, naïve T-cells polarized towards Th17 in the presence of no *B. infantis* fecal water expressed elevated markers of activation and proliferation (Ki67) compared to naïve T-cells polarized towards Th17 in the presence of *B. infantis* EVC001 fecal water (Figure 6E). Collectively, these results imply that during the first weeks of life there are transient immune activation events, centered on the mucosal surfaces, primarily in the gut. The colonization and formation of the gut microbiome plays an integral role in this process and is likely, itself influenced by this process. Specific bacteria, such as *B. infantis* EVC001, influence this process by dampening intestinal inflammation, shaping immune cell-cell regulatory relationships, and polarizing T-cells away from Th2 to Th1-type cells, with potential long-term consequences for immune imprinting and the risk of developing immune-mediated diseases.

## Discussion

It is increasingly clear that early-life gut dysbiosis influences newborn children’s risk of developing autoimmune and allergic diseases later in life; however, the exact mechanisms driving perturbations in the developing immune system have not been fully elucidated. Results here confirm previous findings that highlight a stereotypic postnatal adaptive immune response to environmental exposures, and extend to show an immunological sequence of events, triggered by microbial colonization, resulting in altered patterns of intestinal inflammation. In particular a lack of bifidobacterial early in life is associated with elevated markers of intestinal inflammation and a perturbed immune cell regulatory network.

The transient immune cell expansion events described here are reminiscent of the transient reactions to gut microbes in mice at the time of weaning, termed “the weaning reaction” (Nabhani et al., 2019). It has been postulated that such timed reactivity to the colonizing microbes in mice is regulated by Goblet cell-associated passages (GAPs) and serve important functions in establishing a healthy immune-microbe interface in the gut and the induction of tolerance (Knoop et al., 2017). There are several notable differences between these transient events reported in mice and the process uncovered in human newborns in our current report. First, the timing of transient immune cell reactions is completely different and uncoupled from weaning since the breastfeeding regimen differs between children and most breastfed children in our cohort continue to nurse beyond the three-month window during which the dramatic immune cell changes occur. Secondly, the type of reactions seen in human newborns seem qualitatively distinct and involve different cell populations as compared to the changes seen in mice. This is not only due to the difference between blood-focused analyses in human newborns and intestinal analyses performed in mouse model organisms (Nabhani and Eberl, 2017, 2020; Nabhani et al., 2019), because when analyzing blood immune cell changes in mice, the events remain distinct from human immune cell developmental changes (Brodin lab, manuscript in preparation). The comparison between human newborns and young mice colonized with a more natural gut microbiome will be informative and likely more similar in this regard as shown for other immune system processes (Rosshart et al., 2017).

Furthermore, we report that feeding *B. infantis* EVC001 modulated of immune dysregulation by silencing intestinal Th2 and Th17 immune responses, while inducing IFNβ production, and in vitro its metabolites skewed T-cell polarization *in vitro*, from Th2, towards Th1 phenotype. Thus, given that the gut microbiome seems to play a major role in early immune system development, feeding *B. infantis* to breastfed infants may improve immune imprinting during the first critical months of life. No molecular mechanisms have been identified that directly link microbiome composition or function with the key steps of immune development in humans; however, infants that lack *Bifidobacterium* are more likely to develop autoimmune and asthmatic diseases (Arrieta et al., 2015, 2017; Vatanen et al., 2016). One proposed candidate mechanism involves the over-production of Th17-inducing cytokines leading to increased numbers of Th17 cells as one mechanism of elevated disease risk (Guo et al., 2008; Harrington et al., 2005).

Here we report additional possible mechanisms of immune system imprinting, possibly influencing the risk of developing immune mediated diseases later in life. Our data indicated low abundance of *Bifidobacterium* colonization correlated with increased enteric inflammation both in fecal water and in circulation and is associated with a perturbed cell-cell regulatory network. In particular, memory Tregs frequency was inversely correlated with proinflammatory monocytes and activated T-cell populations in children with abundant *Bifidobacterium*, but this regulatory relationship is lost in children devoid of such beneficial microbes. Indeed, Bifidobacteria are efficient producers of SCFA (Frese et al., 2017), which are in turn important inducers of regulatory T-cells in the gut (Smith et al., 2013). Additionally, we uncovered another possible mechanism of beneficial imprinting by beneficial Bifidobacteria. Breastfed infants supplemented with *B. infantis* EVC001 upregulated intestinal IFNβ, which is known to and promote the development of regulatory T cells (Kotredes et al., 2017). Moreover, IFNβ inhibits Th17 cell differentiation and consequently lowers IL-17 production (Durelli et al., 2009; Ramgolam et al., 2009) and children supplemented with *B. infantis* EVC001 had significantly lower levels of fecal IL-17A as compared to control children. Exogenous IFNβ is given to patients with Multiple Sclerosis (MS) and is believed to improve Treg-mediated immunological tolerance (González-Navajas et al., 2012), inhibit IL-23-dependent Th17 cell expansion (Harrington et al., 2005) and TGB-β/IL-6-mediated Th17 development (Guo et al., 2008). Additionally, there is preclinical evidence of Treg cell mediated protection in a rodent colitis models whereby both intestinal inflammation and cytokine secretion is decreased by exogenous IFNβ (González-Navajas et al., 2012). Collectively these data indicate a previously unappreciated role for IFNβ in early life immune system imprinting by beneficial microbes such as *B. infantis*.

Microbially-derived metabolites are also likely to be critical in immune system development (Lavelle and Sokol, 2020). Indeed, perturbations to the gut microbiome early in life due to antibiotic use lead to changes in the production of bacterial metabolites such as short-chain fatty acids (SCFAs) and other microbial-derived metabolites involved in immune maturation and homeostasis, as well as host energy metabolism, and intestinal integrity (Lavelle and Sokol, 2020). For example, 12-13 DIHOME, a microbial-derived metabolite has recently been shown to correlate with lower risk of asthma development in susceptible infant populations (Levan et al., 2019). Moreover, indole-3-lactic acid (ILA), which is produced in high concentration in breastfed infants colonized with *B. infantis* has been shown to decrease enteric inflammation through activation of Aryl hydrocarbon receptor and NF erythroid 2-related factor (Nrf2) (Ehrlich et al., 2018; Laursen et al., 2020; Meng et al., 2020). Importantly, differences in *B. infantis* HMO utilization loci have been identified (Albert et al., 2019) and strains missing specific genes involved in HMO utilization are unlikely to confer the same benefits to infants as we observed here, as the conversion of indigestible HMOs to infant accessible short chain fatty acids and indole-3-lactic acid are key functions of a healthy gut microbiome (Duar et al., 2020). Similarly, acetate and propionate have been implicated in the reduction of lung inflammation in animal models (Trompette et al., 2014) and development of food allergy in human children (Sandin et al., 2009). Notably, a previous intervention study showed significantly increased SCFAs and lactate production in infants fed *B. infantis* (Frese et al., 2017); due to its unique evolution to consume human milk oligosaccharides (HMOs)(Sela et al., 2008) and a recent study of preterm infants supplemented with *Bifidobacterium* and *Lactobacillus* showed increased concentrations of acetate and lactate as compared to controls not given any microbial supplementation (Alcon-Giner et al., 2020).

Finally, the finding that fecal water from children supplemented with *B. infantis* EVC001 influence T-cell polarization *in vitro* towards Th1 cell states, while fecal water from control children induce Th2-like phenotypes is intriguing and suggest that specific metabolites exert a skewing effect on naïve CD4^+^ T-cells. This effect is probably not mediated by IFNβ alone since cultures supplemented with IFNβ did not show this same pattern (Supplementary Figure 2). Instead, we hypothesize that IFNβ may provide a tolerizing effect in already polarized T-cells. Future metabolomic profiling of the fecal water and functional testing during T-cell polarization, including analyses of epigenetic changes during polarization will be valuable to mechanistically understand this imprinting effect further.

Collectively, the data provided here extends our current understanding of immune system imprinting by beneficial microbes such as *B. infantis* EVC001 and proposes possible explanations for the known differences in risk of developing immune mediated diseases associated with such early life imprinting effects. Randomized controlled trials targeting these additional mediators and mechanisms proposed will be required to establish a way forward towards preventive interventions to lower the burden of immune mediated diseases such as asthma, allergies, and autoimmunity in the future.

## STAR Methods

### Born-Immune newborn cohort study

The study was performed in accordance with the declaration of Helsinki and the study protocol was approved by the regional ethical board in Stockholm, Sweden (DNR: 2009/2052 – 31/3 & 2014/921-32). After obtaining informed consent from parents, blood samples from newborns and parents were collected at the Karolinska University Hospital. We also collected fecal samples from infants, either at the time of clinical visits and frozen directly at −80 C or collected at home frozen at −20 C and brought to the clinic by parents. Clinical metadata such as mode of delivery, nutrition, growth and medications were gathered in a clinical database.

### Blood immune cell profiling by Mass cytometry

Blood samples drawn from newborns and parents were mixed with a stabilizer(Brodin et al., 2019) (one of the components of Whole blood processing kit; Cytodelics AB, Sweden) either immediately or within 1-3 hours post blood draw and cryopreserved as per the manufacturer’s recommendations. Samples were thawed, and cells were fixed/RBCs lysed using WASH # 1 and WASH # 2 buffers (Whole blood processing kit; Cytodelics AB, Sweden) as per the manufacturer’s recommendations. This was performed a few days prior to barcoding and staining of cells. Post fix/lysis of cells, ~1-2×10^6^ cells/sample were plated onto a 96 well round bottom plate using standard cryoprotective solution (10% DMSO and 90% FBS) and cryopreserved at −80°C. At the time of experimentation, cells were thawed at 37°C using RPMI medium supplemented with 10% fetal bovine serum (FBS), 1% penicillin-streptomycin and benzonase (Sigma-Aldrich, Sweden). Briefly, cells were barcoded using automated liquid handling robotic system (Agilent technologies)(Mikes et al., 2019) using the Cell-ID 20-plex Barcoding kit (Fluidigm Inc.) as per the manufacturer’s recommendations. Samples were pooled batch wise by keeping together the longitudinal samples from each new born baby or parent in the same batch. Cells were then washed, FcR blocked using blocking buffer (in-house developed recipe) for 12 min at room temperature, following which cells were incubated for another 30 min at 4°C after addition of a cocktail of metal conjugated antibodies targeting the surface antigens. Cells were washed twice with CyFACS buffer (PBS with 0.1% BSA, 0.05% sodium azide and 2mM EDTA) and fixed overnight using 2% formaldehyde made in PBS (VWR, Sweden). The broad extended panel of antibodies used are listed in Supplementary Table 1. For acquisition by CyTOF, cells were stained with DNA intercalator (0.125 μM Iridium-191/193 or MaxPar^®^ Intercalator-Ir, Fluidigm) in 2% formaldehyde made in PBS for 20 min at room temperature. Cells were washed twice with CyFACS buffer, once with PBS and twice with milliQ water. Cells were mixed with 0.1X Norm Beads (EQ™ Four Element Calibration Beads, Fluidigm) filtered through a 35μm nylon mesh and diluted to 1000,000 cells/ml. Samples were acquired using super samplers connected to our CyTOF2 mass cytometers (Fluidigm Inc.) using CyTOF software version 6.0.626 with noise reduction, a lower convolution threshold of 200, event length limits of 10-150 pushes, a sigma value of 3, and flow rate of 0.045 ml/min.

### Antibodies and reagents for mass cytometry

The panel of monoclonal antibodies used for this study are indicated in the Key Resources Table. Monoclonal antibodies were either purchased pre-conjugated from Fluidigm or obtained in carrier/protein-free buffer as purified antibodies that were then coupled to lanthanide metals using the MaxPar X8 polymer conjugation kit (Fluidigm Inc.) as per the manufacturer’s recommendations. Following the protein concentration determination by measurement of absorbance at 280nm on a nanodrop, the metal-labeled antibodies were diluted in Candor PBS Antibody Stabilization solution (Candor Bioscience, Germany) for long-term storage at 4°C.

### Born-immune Plasma protein profiling

Plasma protein data was generated using Olink assays, a proximity extension assay (Olink AB, Uppsala)(Lundberg et al., 2011). For analysis, 20μL of plasma from each sample was thawed and sent for analysis, either at the plasma protein profiling platform, Science for Life Laboratory, Stockholm or Olink AB in Uppsala. In these assays, plasma proteins are dually recognized by pairs of antibodies coupled to a cDNA-strand that ligates when brought into proximity by its target, extended by a polymerase and detected using a Biomark HD 96.96 dynamic PCR array (Fluidigm Inc.). Three Olink panels have been used as indicated in Key Resources Table, capturing a total of 267 unique proteins in each plasma sample.

### Born-immune fecal metagenomics

DNA was extracted as in IHMS DNA extraction protocol #8 (Costea et al., 2017). According to protocol recommendations 0,2g of faeces were used. Briefly, samples were treated with lysozyme solution and subjected to bead beating using zirconium beads. After centrifugation, DNA is extracted from supernatants with QIAamp Fast DNA Stool Mini Kit (Qiagen, Cat No. 51604). After DNA extraction, collected DNA was quantified with Qubit (ThermoFisher, Cat No. Q32851) and 10ng were subjected to mechanical fragmentation with the Covaris Focused-ultrasonicator to ensure that fragment sizes were compatible with Illumina sequencing (~300bp average). Sequencing adapters and sample barcodes were incorporated to the DNA fragments using ThruPLEX DNA-seq kit (Rubicon Genomics, Cat No. R400406). ThruPLEX DNA-seq products were purified and size selected by AMPure beads (Beckman Coulter, Cat No. B23318), and DNA concentration and size distribution were inspected with the Qubit dsDNA HS Assay Kit (ThermoFisher, Cat No. Q32851) and the Agilent 2100 Bioanalyzer High Sensitivity kit (Agilent Technologies, Cat No. 5067-4626), respectively. Purified ThruPLEX DNA-seq products were then equimolarly pooled in 4 lanes and subjected to NovaSeq 6000 S4 Illumina Sequencing at the Science for Life Laboratory, Stockholm, Sweden.

### IMPRINT study design and cytokine quantification

A total of 80 fecal samples (ClinicalTrials.gov: NCT02457338) from 40 subjects were analyzed at three time points—day 6 (Baseline), day 40, and day 60 postnatally. Individual subjects were chosen at random and made up a subset of the original study participants. All aspects of the study were approved by the University of California Davis Institutional Review Board (IRB Number: ID 631099) and all participants provided written informed consent. Details of the study design and procedures used to collect these samples has been reported (Frese et al., 2017). Briefly, exclusively breastfed term infants were randomly selected to receive 1.8 × 10^10^ colony-forming units (CFU) B. infantis EVC001 daily for 21 days (EVC001) starting at day 7 postnatal or to receive breast milk alone (control) and followed up to postnatal day 60 (Frese et al., 2017). All mothers received lactation support throughout the study. The demographic information (e.g., age, sex, and gestational age) was collected from each participant. Here stool samples from randomly selected infants who were fed EVC001 (n = 20) and control infants (n = 20) on days 6 (Baseline) and 60 postnatal were collected and quantified from 80 mg of frozen stool diluted 1:10 in Meso Scale Discovery (MSD; Rockville, MD) diluent using the U-PLEX Inflammation Panel 1 (human) Kit. Samples were measured in duplicate and blank values were subtracted from all readings according to the manufacturer’s instructions. All detectable biomarker values were included as continuous data in the analyses; however, values below level of detection (< 30% of all cytokines measurements) were generated below the level of quantification to justify parametric statistics. Fecal cytokine concentrations were determined using calibrations curves to which electrochemiluminescence signals were backfitted. Final concentrations were calculated using the Sector Imager 2400 MSD Discovery Workbench analysis software.

### IMPRINT study – fecal microbiome analyses

DNA was extracted from 296 stool swab samples stored in DNA/RNA shield lysis tubes (Zymo Research, Irvine CA) using the ZymoBIOMICS 96 MagBead DNA kit (Zymo Research). Extracted DNA was quantified using QuantIT dsDNA Assay kit, high sensitivity (ThermoFisher Scientific, Waltham, MA) according to the manufacturer’s protocol. 3 samples were omitted from downstream analysis due to failure to meet input requirements for library preparation. Libraries were prepared for each sample using the Illumina Nextera DNA Flex library kit (Illumina, San Diego, CA) with unique dual indexes according to manufacturer guidelines. Libraries were pooled and submitted to UC Davis DNA Technologies core for sequencing on the Illumina NovaSeq S4 flow cell. (Illumina, San Diego, CA). Each lane of the S4 flow cell contained 96 libraries.

### IMPRINT-Absolute quantification of *B. infantis* by Quantitative Real-Time PCR

Quantification of the total *B. infantis* was performed by quantitative real-time PCR using Blon_2348 sialidasetgene primers Inf2348F (5’- ATA CAG CAG AAC CTT GGC CT -3’), Inf2348_R (5’- GCG ATC ACA TGG ACG AGA AC -3’), and Inf2348_P (5’- /56-FAM/TTT CAC GGA /ZEN/TCA CCG GAC CAT ACG /3lABkFQ/ -3’)(Lawley et al., 2017) Each reaction contained 10μL of 2× TaqMan Universal Master Mix II with UNG master mix (Applied Biosystems), 0.9 μm of each primer, 0.25 μM probe and 5 μL of template DNA. Thermal cycling was performed on a QuantStudio 3 Real-Time PCR System and consisted of an initial UNG activation step step of 2 minute at 50 followed by a 10 minute denaturation at 95°Csucceeded by 40 cycles of 15 s at 95 °C and 1 min at 60°C. Standard curves for absolute quantification were generated using genomic DNA extracted from a pure culture of *B. infantis* EVC001.

### IMPRINT-Quality filtering and removal of human sequences

Demultiplexed fastq sequences were quality filtered, including adaptor trimming using Trimmomatic v0.36 (Czajkowski et al., 2018) with default parameters. Quality-filtered sequences were screened to remove human sequences using GenCoF v1.0 (Czajkowski et al., 2018) against a non-redundant version of the Genome Reference Consortium Human Build 38, patch release 7 (GRCh38_p7; www.ncbi.nlm.nih.gov). Human sequence-filtered raw reads were deposited in the Sequence Read Archive (SRA; https://www.ncbi.nlm.nih.gov/sra) under the reference number, PRJNA390646.

### IMPRINT-Fecal water preparation

Historical fecal samples from term infants either colonized with *B. infantis* EVC001 or not were collected and stored in −80C until processing. Pooled fecal samples (minimum of 3) from infants colonized with *B. infantis* EVC001 or not were weighed and diluted 25% w/v in sterile PBS and vortexed for 1 min allowing stool to thaw and form a homogenous slurry. Fecal slurries were then centrifuged for 30min at 4,000 RPM at 4C. The supernatant was collected and spun again for 3 hours at 12,000 RPM at 4C. The supernatant was further collected and then serially filtered (40um cell strainer, 1um, .45um, and .22um). Filtered fecal waters were stored in −80C until use.

### IMPRINT-fecal cytokine measurements

Interleukin (IL)-1 β, IL-2, IL-4 IL-5, IL-8, IL-10, IL-13, IL-17A, IFNγ, and TNFα were quantified from 80 mg of stool diluted 1:10 in Meso Scale Discovery (MSD; Rockville, MD) diluent using the U-PLEX Inflammation Panel 1 (human) Kit according to the manufacturer’s instructions as previously published (ref). Standards and samples were measured in duplicate and blank values were subtracted from all readings. The concentration of fecal calprotectin and MPO were quantified using MSD R-PLEX Human Calprotectin Antibody Set. The samples were plated in duplicate and the assay was performed twice. The plates were then read on a Sector Imager 2400 MSD Discovery Workbench analysis software.

### IMPRINT - Statistical Analysis

All statistical analyses were performed in R v3.6.2. A Kruskal-Wallis one-way analysis of variance coupled with an FDR or Bonferroni correction was used for statistical comparisons between individual genes, cytokines and taxa amongst groups. Statistical analysis to assess total resistome or enterotype composition by group was performed using a Mann-Whitney or Holm-adjusted Dunn’s test. Rarefaction curves were computed to estimate the diversity of the identified ARGs across samples. A nonparametric two-sample t-test was used to compare rarefaction curves using Monte Carlo permutations (n=999). Enterotype analysis was performed as previously described (Arumugam et al., 2011). Cytokines and *B. infantis-specific* qPCR were correlated with the Spearman method with FDR correction. The P-values throughout the manuscript are represented by asterisks (*, *P* < 0.05; **, *P* < 0.01; ***, *P* < 0.001; ****, *P* < 0.0001). A random forest model was created to identify taxa most often used in prediction of IFN concentration. Family-level taxa relative abundance at days 40 and 60 and age (day of life) were used as predictors. The depth at which each taxon appeared within a tree was collected and averaged across the ensemble of 500 trees.

## ACKNOWLEDGEMENTS

The authors thank the mothers and their infants enrolled in the clinical trial for collecting information and samples with methodological detail.

## AUTHOR CONTRIBUTIONS

BMH and PB conceived the study. BMH, JTS, MAU, JBG, SAF, and PB designed the study. RDM, SC, JP, and HKB generated data presented in the manuscript. BMH, JTS, JBG, SAF, and PB analyzed the data. BMH, AME, and PB wrote the manuscript. All authors approved the final manuscript for publication.

---

**Supplementary Figure 1.**
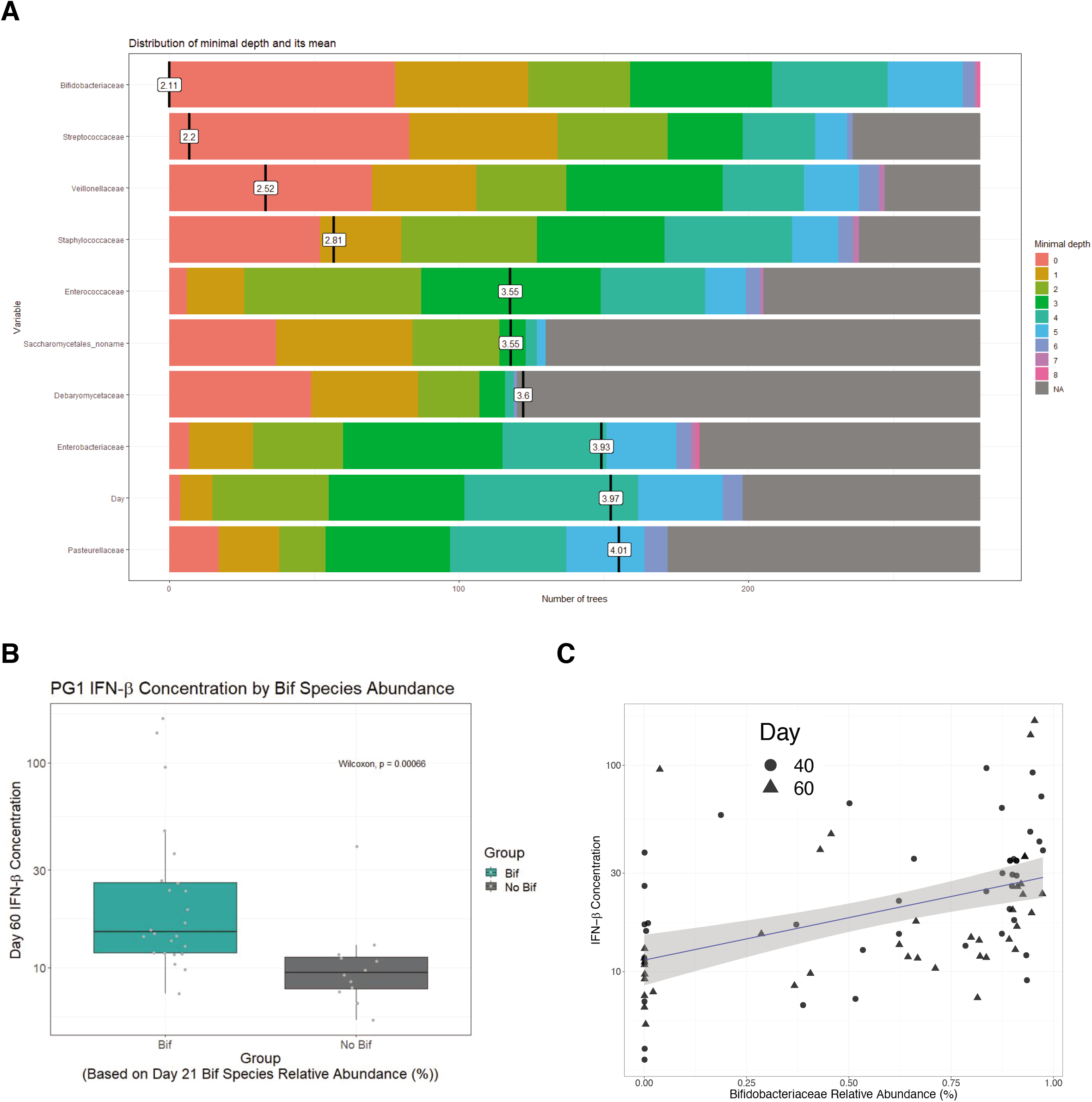
Related to Figure 5. Gut microbes associated with fecal cytokine levels. (**A**) Random Forest Model output. (**B**) Box plot of day 60 fecal IFNb abundances. (**C**) Correlation (Pearson) between day 60 relative abundance of Bifidobacteria and fecal IFNb, r=0.55, p=8.9 x 10^-8^

**Supplementary Figure 2.**
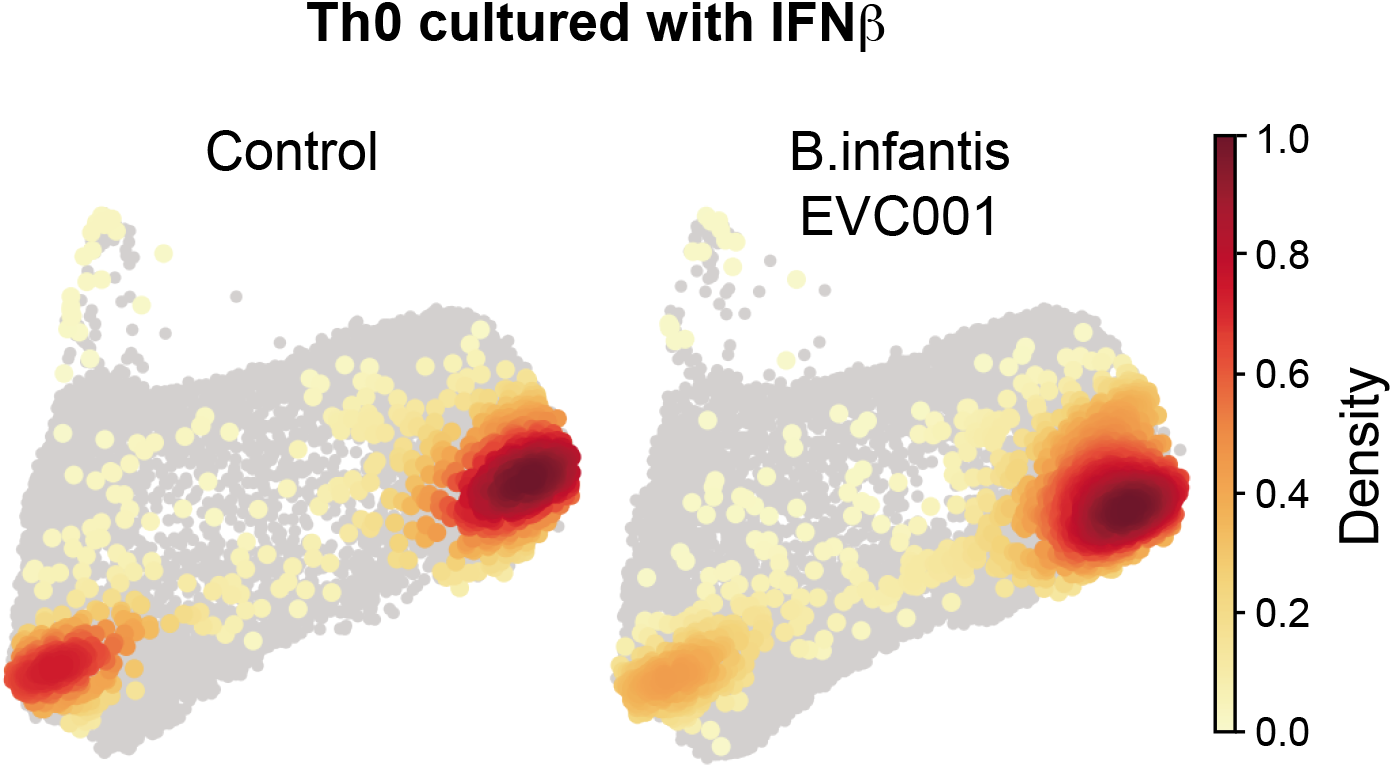
Related to Figure 6. CD4^+^ T-cell polarization under the influence of microbial metabolites and IFNβ. CD4^+^ T-cell polarization in vitro in the presence of fecal water from children given B.infantis EVC001 or control and additional IFNβ. T-cells at the end of the culture shown as 2D PAGA plots and separated by culture condition.

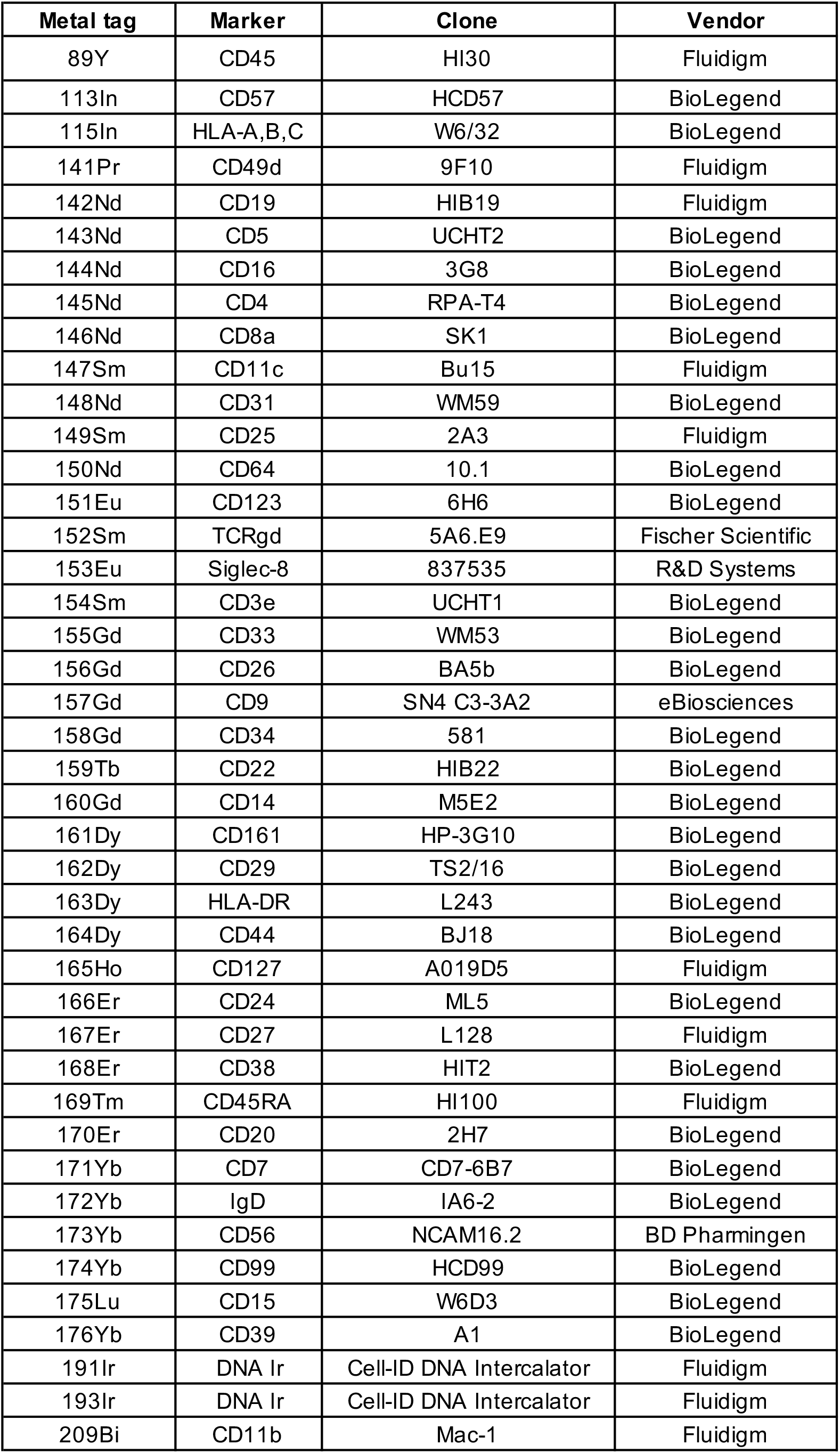

| **Metal tag** | **Marker** | **Clone** | **Vendor** |
| --- | --- | --- | --- |
| 89Y | CD45 | HI30 | Fluidigm |
| 113In | CD57 | HCD57 | BioLegend |
| 115In | HLA-A,B,C | W6/32 | BioLegend |
| 141Pr | CD49d | 9F10 | Fluidigm |
| 142Nd | CD19 | HIB19 | Fluidigm |
| 143Nd | CD5 | UCHT2 | BioLegend |
| 144Nd | CD16 | 3G8 | BioLegend |
| 145Nd | CD4 | RPA-T4 | BioLegend |
| 146Nd | CD8a | SK1 | BioLegend |
| 147Sm | CD11c | Bu15 | Fluidigm |
| 148Nd | CD31 | WM59 | BioLegend |
| 149Sm | CD25 | 2A3 | Fluidigm |
| 150Nd | CD64 | 10.1 | BioLegend |
| 151Eu | CD123 | 6H6 | BioLegend |
| 152Sm | TCRgd | 5A6.E9 | Fischer Scientific |
| 153Eu | Siglec-8 | 837535 | R&D Systems |
| 154Sm | CD3e | UCHT1 | BioLegend |
| 155Gd | CD33 | WM53 | BioLegend |
| 156Gd | CD26 | BA5b | BioLegend |
| 157Gd | CD9 | SN4 C3-3A2 | eBiosciences |
| 158Gd | CD34 | 581 | BioLegend |
| 159Tb | CD22 | HIB22 | BioLegend |
| 160Gd | CD14 | M5E2 | BioLegend |
| 161Dy | CD161 | HP-3G10 | BioLegend |
| 162Dy | CD29 | TS2/16 | BioLegend |
| 163Dy | HLA-DR | L243 | BioLegend |
| 164Dy | CD44 | BJ18 | BioLegend |
| 165Ho | CD127 | A019D5 | Fluidigm |
| 166Er | CD24 | ML5 | BioLegend |
| 167Er | CD27 | L128 | Fluidigm |
| 168Er | CD38 | HIT2 | BioLegend |
| 169Tm | CD45RA | HI100 | Fluidigm |
| 170Er | CD20 | 2H7 | BioLegend |
| 171Yb | CD7 | CD7-6B7 | BioLegend |
| 172Yb | IgD | IA6-2 | BioLegend |
| 173Yb | CD56 | NCAM16.2 | BD Pharmingen |
| 174Yb | CD99 | HCD99 | BioLegend |
| 175Lu | CD15 | W6D3 | BioLegend |
| 176Yb | CD39 | A1 | BioLegend |
| 191Ir | DNA Ir | Cell-ID DNA Intercalator | Fluidigm |
| 193Ir | DNA Ir | Cell-ID DNA Intercalator | Fluidigm |
| 209Bi | CD11b | Mac-1 | Fluidigm |

## REFERENCES

Abrahamsson, T.R., Jakobsson, H.E., Andersson, A.F., Björkstén, B., Engstrand, L., and Jenmalm, M.C. (2014). Low gut microbiota diversity in early infancy precedes asthma at school age. Clin Exp Allergy 44, 842–850.

Albert, K., Rani, A., and Sela, D.A. (2019). Comparative Pangenomics of the Mammalian Gut Commensal Bifidobacterium longum. Microorg 8, 7.

Alcon-Giner, C., Dalby, M.J., Caim, S., Ketskemety, J., Shaw, A., Sim, K., Lawson, M.A.E., Kiu, R., Leclaire, C., Chalklen, L., et al. (2020). Microbiota Supplementation with Bifidobacterium and Lactobacillus Modifies the Preterm Infant Gut Microbiota and Metabolome: An Observational Study. Cell Reports Medicine 1, 100077.

Arrieta, M.-C., Stiemsma, L.T., Dimitriu, P.A., Thorson, L., Russell, S., Yurist-Doutsch, S., Kuzeljevic, B., Gold, M.J., Britton, H.M., Lefebvre, D.L., et al. (2015). Early infancy microbial and metabolic alterations affect risk of childhood asthma. Science Translational Medicine 7, 307ra152–307ra152.

Arrieta, M.-C., Arévalo, A., Stiemsma, L., Dimitriu, P., Chico, M.E., Loor, S., Vaca, M., Boutin, R., Morien, E., Jin, M., et al. (2017). Associations between infant fungal and bacterial dysbiosis and childhood atopic wheeze in a nonindustrialized setting. Journal of Allergy and Clinical Immunology.

Arumugam, M., Raes, J., Pelletier, E., Paslier, D.L., Yamada, T., Mende, D.R., Fernandes, G.R., Tap, J., Bruls, T., Batto, J.-M., et al. (2011). Enterotypes of the human gut microbiome. Nature 473, 174–180.

Avershina, E., Storrø, O., Øien, T., Johnsen, R., Pope, P., and Rudi, K. (2014). Major faecal microbiota shifts in composition and diversity with age in a geographically restricted cohort of mothers and their children. Fems Microbiol Ecol 87, 280–290.

Azad, M.B., Konya, T., Maughan, H., Guttman, D.S., Field, C.J., Chari, R.S., Sears, M.R., Becker, A.B., Scott, J.A., Kozyrskyj, A.L., et al. (2013). Gut microbiota of healthy Canadian infants: profiles by mode of delivery and infant diet at 4 months. Can Med Assoc J 185, 385–394.

Brodin, P. (2020). New approaches to the study of immune responses in humans. Hum Genet 1–5.

Brodin, P., Duffy, D., and Quintana-Murci, L. (2019). A Call for Blood—In Human Immunology. Immunity 50, 1335–1336.

Cano-Gamez, E., Soskic, B., Roumeliotis, T.I., So, E., Smyth, D.J., Baldrighi, M., Willé, D., Nakic, N., Esparza-Gordillo, J., Larminie, C.G.C., et al. (2020). Single-cell transcriptomics identifies an effectorness gradient shaping the response of CD4+ T cells to cytokines. Nat Commun 11, 1801.

Costea, P.I., Zeller, G., Sunagawa, S., Pelletier, E., Alberti, A., Levenez, F., Tramontano, M., Driessen, M., Hercog, R., Jung, F.-E., et al. (2017). Towards standards for human fecal sample processing in metagenomic studies. Nat Biotechnol 35, 1069–1076.

Czajkowski, M.D., Vance, D.P., Frese, S.A., and Casaburi, G. (2018). GenCoF: a graphical user interface to rapidly remove human genome contaminants from metagenomic datasets. Bioinformatics 35, 2318–2319.

Davis, M.M., and Brodin, P. (2018). Rebooting Human Immunology. Annu Rev Immunol 36, 1–22.

Dominguez-Bello, M., Godoy-Vitorino, F., Knight, R., and Blaser, M.J. (2019). Role of the microbiome in human development. Gut 68, 1108.

Duar, R.M., Henrick, B.M., Casaburi, G., and Frese, S.A. (2020). Integrating the Ecosystem Services Framework to Define Dysbiosis of the Breastfed Infant Gut: The Role of B. infantis and Human Milk Oligosaccharides. Frontiers Nutrition 7, 33.

Durelli, L., Conti, L., Clerico, M., Boselli, D., Contessa, G., Ripellino, P., Ferrero, B., Eid, P., and Novelli, F. (2009). T-helper 17 cells expand in multiple sclerosis and are inhibited by interferon-β. Ann Neurol 65, 499–509.

Ehrlich, A.M., Henrick, B., Pacheco, A., Taft, D., Xu, G., Huda, N., Lozada-Contreras, M., Goodson, M., Slupsky, C., Mills, D., et al. (2018). Bifidobacterium grown on human milk oligosaccharides produce tryptophan metabolite Indole-3-lactic acid that significantly decreases inflammation in intestinal cells in vitro. Faseb J 32, lb359–lb359.

Frese, S.A., Hutton, A.A., Contreras, L.N., Shaw, C.A., Palumbo, M.C., Casaburi, G., Xu, G., Davis, J.C.C., Lebrilla, C.B., Henrick, B.M., et al. (2017). Persistence of Supplemented Bifidobacterium longum subsp. infantis EVC001 in Breastfed Infants. Msphere 2, e00501–17.

González-Navajas, J.M., Lee, J., David, M., and Raz, E. (2012). Immunomodulatory functions of type I interferons. Nat Rev Immunol 12, 125–135.

Grzekowiak, ukasz, Collado, M.C., Mangani, C., Maleta, K., Laitinen, K., Ashorn, P., Isolauri, E., and Salminen, S. (2012). Distinct Gut Microbiota in Southeastern African and Northern European Infants. J Pediatr Gastr Nutr 54, 812–816.

Guo, B., Chang, E.Y., and Cheng, G. (2008). The type I IFN induction pathway constrains Th17-mediated autoimmune inflammation in mice. J Clin Invest 118, 1680–1690.

Harrington, L.E., Hatton, R.D., Mangan, P.R., Turner, H., Murphy, T.L., Murphy, K.M., and Weaver, C.T. (2005). Interleukin 17–producing CD4+ effector T cells develop via a lineage distinct from the T helper type 1 and 2 lineages. Nat Immunol 6, 1123–1132.

Henrick, B.M., Chew, S., Casaburi, G., Brown, H.K., Frese, S.A., Zhou, Y., Underwood, M.A., and Smilowitz, J.T. (2019). Colonization by B. infantis EVC001 modulates enteric inflammation in exclusively breastfed infants. Pediatr Res 86, 749–757.

Huda, M.N., Lewis, Z., Kalanetra, K.M., Rashid, M., Ahmad, S.M., Raqib, R., Qadri, F., Underwood, M.A., Mills, D.A., and Stephensen, C.B. (2014). Stool Microbiota and Vaccine Responses of Infants. Pediatrics 134, e362–e372.

Hviid, A., Svanström, H., and Frisch, M. (2011). Antibiotic use and inflammatory bowel diseases in childhood. Gut 60, 49.

Ioannidis, M., Cerundolo, V., and Salio, M. (2020). The Immune Modulating Properties of Mucosal-Associated Invariant T Cells. Front Immunol 11, 1556.

Jost, T., Lacroix, C., Braegger, C.P., and Chassard, C. (2012). New Insights in Gut Microbiota Establishment in Healthy Breast Fed Neonates. Plos One 7, e44595.

Knoop, K.A., Gustafsson, J.K., McDonald, K.G., Kulkarni, D.H., Coughlin, P.E., McCrate, S., Kim, D., Hsieh, C.-S., Hogan, S.P., Elson, C.O., et al. (2017). Microbial antigen encounter during a preweaning interval is critical for tolerance to gut bacteria. Science Immunology 2, eaao1314.

Kotredes, K.P., Thomas, B., and Gamero, A.M. (2017). The Protective Role of Type I Interferons in the Gastrointestinal Tract. Front Immunol 8, 410.

Laforest-Lapointe, I., and Arrieta, M.-C. (2017). Patterns of Early-Life Gut Microbial Colonization during Human Immune Development: An Ecological Perspective. Front Immunol 8, 788.

Laursen, M.F., Sakanaka, M., Burg, N. von, Andersen, D., Mörbe, U., Rivollier, A., Pekmez, C.T., Moll, J.M., Michaelsen, K.F., Mølgaard, C., et al. (2020). Breastmilk-promoted bifidobacteria produce aromatic lactic acids in the infant gut. Biorxiv 2020.01.22.914994.

Lavelle, A., and Sokol, H. (2020). Gut microbiota-derived metabolites as key actors in inflammatory bowel disease. Nat Rev Gastroentero 17, 223–237.

Lawley, B., Munro, K., Hughes, A., Hodgkinson, A.J., Prosser, C.G., Lowry, D., Zhou, S.J., Makrides, M., Gibson, R.A., Lay, C., et al. (2017). Differentiation of Bifidobacterium longum subspecies longum and infantis by quantitative PCR using functional gene targets. Peerj 5, e3375.

Levan, S.R., Stamnes, K.A., Lin, D.L., Panzer, A.R., Fukui, E., McCauley, K., Fujimura, K.E., McKean, M., Ownby, D.R., Zoratti, E.M., et al. (2019). Elevated faecal 12,13-diHOME concentration in neonates at high risk for asthma is produced by gut bacteria and impedes immune tolerance. Nat Microbiol 4, 1851–1861.

Lewis, Z.T., Totten, S.M., Smilowitz, J.T., Popovic, M., Parker, E., Lemay, D.G., Tassell, M.L.V., Miller, M.J., Jin, Y.-S., German, J.B., et al. (2015). Maternal fucosyltransferase 2 status affects the gut bifidobacterial communities of breastfed infants. Microbiome 3, 13.

Li, S., Rouphael, N., Duraisingham, S., Romero-Steiner, S., Presnell, S., Davis, C., Schmidt, D.S., Johnson, S.E., Milton, A., Rajam, G., et al. (2013). Molecular signatures of antibody responses derived from a systems biology study of five human vaccines. Nature Immunology 15, 195–204.

LoCascio, R.G., Desai, P., Sela, D.A., Weimer, B., and Mills, D.A. (2010). Broad Conservation of Milk Utilization Genes in Bifidobacterium longum subsp. infantis as Revealed by Comparative Genomic Hybridization ▿ †. Appl Environ Microb 76, 7373–7381.

Lundberg, M., Eriksson, A., Tran, B., Assarsson, E., and Fredriksson, S. (2011). Homogeneous antibody-based proximity extension assays provide sensitive and specific detection of low-abundant proteins in human blood. Nucleic Acids Res 39, e102–e102.

Maggi, L., Santarlasci, V., Capone, M., Peired, A., Frosali, F., Crome, S.Q., Querci, V., Fambrini, M., Liotta, F., Levings, M.K., et al. (2010). CD161 is a marker of all human IL-17-producing T-cell subsets and is induced by RORC. Eur J Immunol 40, 2174–2181.

Mair, F., Erickson, J.R., Voillet, V., Simoni, Y., Bi, T., Tyznik, A.J., Martin, J., Gottardo, R., Newell, E.W., and Prlic, M. (2020). A Targeted Multi-omic Analysis Approach Measures Protein Expression and Low-Abundance Transcripts on the Single-Cell Level. Cell Reports 31, 107499.

Meng, D., Sommella, E., Salviati, E., Campiglia, P., Ganguli, K., Djebali, K., Zhu, W., and Walker, W.A. (2020). Indole-3-lactic acid, a metabolite of tryptophan, secreted by Bifidobacterium longum subspecies infantis is anti-inflammatory in the immature intestine. Pediatr Res 88, 209–217.

Mikes, J., Olin, A., Lakshmikanth, T., Chen, Y., and Brodin, P. (2019). Mass Cytometry, Methods and Protocols. Methods Mol Biology Clifton N J 1989, 111–123.

Mohammadkhah, A.I., Simpson, E.B., Patterson, S.G., and Ferguson, J.F. (2018). Development of the Gut Microbiome in Children, and Lifetime Implications for Obesity and Cardiometabolic Disease. Children 5, 160.

Nabhani, Z., and Eberl, G. (2017). GAPs in early life facilitate immune tolerance. Science Immunology 2, eaar2465.

Nabhani, Z.A., and Eberl, G. (2020). Imprinting of the immune system by the microbiota early in life. Mucosal Immunol 13, 183–189.

Nabhani, Z., Dulauroy, S., Marques, R., Cousu, C., Bounny, S., Déjardin, F., Sparwasser, T., Bérard, M., Cerf-Bensussan, N., and Eberl, G. (2019). A Weaning Reaction to Microbiota Is Required for Resistance to Immunopathologies in the Adult. Immunity.

Olin, A., Henckel, E., Chen, Y., Lakshmikanth, T., Pou, C., Mikes, J., Gustafsson, A., Bernhardsson, A.K., Zhang, C., Bohlin, K., et al. (2018). Stereotypic Immune System Development in Newborn Children. Cell 174, 1277–1292.e14.

Pou, C., Nkulikiyimfura, D., Henckel, E., Olin, A., Lakshmikanth, T., Mikes, J., Wang, J., Chen, Y., Bernhardsson, A., Gustafsson, A., et al. (2019). The repertoire of maternal anti-viral antibodies in human newborns. Nature Medicine 1–6.

Pré, F.M. du, Berkel, L.A. van, Ráki, M., Leeuwen, M.A. van, Ruiter, L.F. de, Broere, F., Borg, M.N.D. ter, Lund, F.E., Escher, J.C., Lundin, K.E.A., et al. (2011). CD62LnegCD38+ Expression on Circulating CD4+ T Cells Identifies Mucosally Differentiated Cells in Protein Fed Mice and in Human Celiac Disease Patients and Controls. Am J Gastroenterol 106, 1147–1159.

Ramgolam, V.S., Sha, Y., Jin, J., Zhang, X., and Markovic-Plese, S. (2009). IFN-β Inhibits Human Th17 Cell Differentiation. J Immunol 183, 5418–5427.

Renz, H., and Skevaki, C. (2020). Early life microbial exposures and allergy risks: opportunities for prevention. Nat Rev Immunol 1–15.

Rhoads, J.M., Collins, J., Fatheree, N.Y., Hashmi, S.S., Taylor, C.M., Luo, M., Hoang, T.K., Gleason, W.A., Arsdall, M.R.V., Navarro, F., et al. (2018). Infant Colic Represents Gut Inflammation and Dysbiosis. J Pediatrics 203, 55–61.e3.

Rodriguez, L., Pekkarinen, P.T., Lakshmikanth, T., Tan, Z., Consiglio, C.R., Pou, C., Chen, Y., Mugabo, C.H., Nguyen, N.A., Nowlan, K., et al. (2020). Systems-level immunomonitoring from acute to recovery phase of severe COVID-19. Cell Reports Medicine 100078.

Roos, S., Dicksved, J., Tarasco, V., Locatelli, E., Ricceri, F., Grandin, U., and Savino, F. (2013). 454 Pyrosequencing Analysis on Faecal Samples from a Randomized DBPC Trial of Colicky Infants Treated with Lactobacillus reuteri DSM 17938. Plos One 8, e56710.

Rosshart, S.P., Vassallo, B.G., Angeletti, D., Hutchinson, D.S., Morgan, A.P., Takeda, K., Hickman, H.D., McCulloch, J.A., Badger, J.H., Ajami, N.J., et al. (2017). Wild Mouse Gut Microbiota Promotes Host Fitness and Improves Disease Resistance. Cell 171, 1015–1028.e13.

Sandin, A., Bråbäck, L., Norin, E., and Björkstén, B. (2009). Faecal short chain fatty acid pattern and allergy in early childhood. Acta Paediatr 98, 823–827.

Sela, D.A., and Mills, D.A. (2010). Nursing our microbiota: molecular linkages between bifidobacteria and milk oligosaccharides. Trends Microbiol 18, 298–307.

Sela, D.A., Chapman, J., Adeuya, A., Kim, J.H., Chen, F., Whitehead, T.R., Lapidus, A., Rokhsar, D.S., Lebrilla, C.B., German, J.B., et al. (2008). The genome sequence of Bifidobacterium longum subsp. infantis reveals adaptations for milk utilization within the infant microbiome. Proc National Acad Sci 105, 18964–18969.

Shao, Y., Forster, S.C., Tsaliki, E., Vervier, K., Strang, A., Simpson, N., Kumar, N., Stares, M.D., Rodger, A., Brocklehurst, P., et al. (2019). Stunted microbiota and opportunistic pathogen colonization in caesarean-section birth. Nature 574, 117–121.

Shin, N.-R., Whon, T.W., and Bae, J.-W. (2015). Proteobacteria: microbial signature of dysbiosis in gut microbiota. Trends Biotechnol 33, 496–503.

Smith, P.M., Howitt, M.R., Panikov, N., Michaud, M., Gallini, C.A., Bohlooly-Y, M., Glickman, J.N., and Garrett, W.S. (2013). The Microbial Metabolites, Short-Chain Fatty Acids, Regulate Colonic T_reg_ Cell Homeostasis. Science 341, 569–573.

Sonnenburg, J.L., and Sonnenburg, E.D. (2019). Vulnerability of the industrialized microbiota. Science 366, eaaw9255.

Trompette, A., Gollwitzer, E.S., Yadava, K., Sichelstiel, A.K., Sprenger, N., Ngom-Bru, C., Blanchard, C., Junt, T., Nicod, L.P., Harris, N.L., et al. (2014). Gut microbiota metabolism of dietary fiber influences allergic airway disease and hematopoiesis. Nat Med 20, 159–166.

Underwood, M.A., German, J.B., Lebrilla, C.B., and Mills, D.A. (2015). Bifidobacterium longum subspecies infantis: champion colonizer of the infant gut. Pediatr Res 77, 229–235.

Vatanen, T., Kostic, A.D., d’Hennezel, E., Siljander, H., Franzosa, E.A., Yassour, M., Kolde, R., Vlamakis, H., Arthur, T.D., Hämäläinen, A.-M., et al. (2016). Variation in Microbiome LPS Immunogenicity Contributes to Autoimmunity in Humans. Cell 165, 842–853.

West, N.R., Hegazy, A.N., Owens, B.M.J., Bullers, S.J., Linggi, B., Buonocore, S., Coccia, M., Görtz, D., This, S., Stockenhuber, K., et al. (2017). Oncostatin M drives intestinal inflammation and predicts response to tumor necrosis factor–neutralizing therapy in patients with inflammatory bowel disease. Nat Med 23, 579–589.

Wolf, F.A., Hamey, F.K., Plass, M., Solana, J., Dahlin, J.S., Göttgens, B., Rajewsky, N., Simon, L., and Theis, F.J. (2019). PAGA: graph abstraction reconciles clustering with trajectory inference through a topology preserving map of single cells. Genome Biol 20, 59.

